# PD1+ T-cells correlate with Nerve Fiber Density as a prognostic biomarker in patients with resected perihilar cholangiocarcinoma

**DOI:** 10.1101/2022.01.07.475344

**Authors:** Xiuxiang Tan, Mika Rosin, Simone Appinger, Jan Bednarsch, Dong Liu, Georg Wiltberger, Juan Garcia Vallejo, Sven Lang, Zoltan Czigany, Shiva Boroojerdi, Nadine T. Gaisa, Peter Boor, Roman David Bülow, Judith de Vos-Geelen, Liselot Valkenburg-van Iersel, Marian Clahsen-van Groningen, Evelien J.M. de Jong, Bas Groot Koerkamp, Michail Doukas, Flavio G. Rocha, Tom Luedde, Uwe Klinge, Shivan Sivakumar, Ulf Neumann, Lara Heij

**Affiliations:** Department of Surgery and Transplantation, University Hospital RWTH Aachen, Aachen, Germany; NUTRIM School of Nutrition and Translational Research in Metabolism, Maastricht University, Maastricht, The Netherlands; Department of Molecular Cell Biology & Immunology, VU University Medical Center, Amsterdam, The Netherlands; Institute of Pathology, University Hospital RWTH Aachen, Aachen, Germany; Department of Internal Medicine, Division of Medical Oncology, GROW School for Oncology and Developmental Biology, Maastricht UMC+, Maastricht, the Netherlands; Department of Pathology, Erasmus University Medical Center, Rotterdam, the Netherlands; Department of Surgery, Erasmus MC Cancer Institute, Rotterdam, the Netherlands; Division of Surgical Oncology, Knight Cancer Institute, Oregon Health and Science University, Portland, OR, USA; Department of Gastroenterology, Hepatology and Infectious Diseases, University Hospital Duesseldorf, Düsseldorf, Germany; Kennedy Institute of Rheumatology, University of Oxford, Oxford, UK; Department of Oncology, University of Oxford, Oxford, UK; Department of Surgery, Maastricht University Medical Centre (MUMC), Maastricht, Netherlands

**Keywords:** Cholangiocarcinoma, Liver cancer, Tumor microenvironment, Nerve fiber density, Immune checkpoint, Immune cells

## Abstract

**Background & Aims:** Perihilar cholangiocarcinoma (pCCA) is a rare hepatobiliary malignancy. Nerve fiber invasion (NFI) shows cancer invading the nerve and is considered an aggressive feature. Nerve fiber density (NFD) consists of small nerve fibers without cancer invasion and is divided into high NFD (high numbers of small nerve fibers) or low NFD (low numbers of small nerve fibers). We aim to explore differences in immune cell populations and survival.

**Methods:** We applied multiplex immunofluorescence (mIF) on 47 pCCA surgically resected patients and investigated the immune cell composition in the tumor microenvironment (TME) of different nerve fiber phenotypes (NFI, high and low NFD). Extensive group comparison was carried out and the association with overall survival (OS) was assessed.

**Results:** The NFI ROI was measured with highest CD68+ macrophage levels among 3 ROIs (NFI compared to tumor free p= 0.016 and to tumor p=0.034). Further, for NFI patients the density of co-inhibitory markers CD8+PD1+ and CD68+PD1+ were more abundant in the tumor rather than NFI ROI (p= 0.004 and p= 0.0029 respectively). Comparison between patients with NFD and NFI groups, the signals of co-expression of CD8+PD1+ as well as CD68+PD1+ were significantly higher in the high NFD group (p= 0.027 and p= 0.044, respectively). The OS for high NFD patients was 92 months median OS (95% CI:41-142), for low NFD patients 20 months ((95% CI: 4-36) and for NFI group of patients 19 months (95% CI 7-33). The OS for high NFD patients was significantly better compared to low NFD (p= 0.046) and NFI (p= 0.032).

**Conclusions:** PD1+ T-cells correlate with high NFD as a prognostic biomarker, the biological pathway behind this needs to be investigated.

**Lay summary:** Nerve fibers play a dual role in the tumor microenvironment in pCCA. In our previous study, we showed that the presence of high numbers of small nerve fibers is associated with a better overall survival. In addition, we found that in high NFD patients PD1+ T-cells are significantly overexpressed. Therefore, we present high NFD as a promising prognostic biomarker.

**Graphical abstract:** 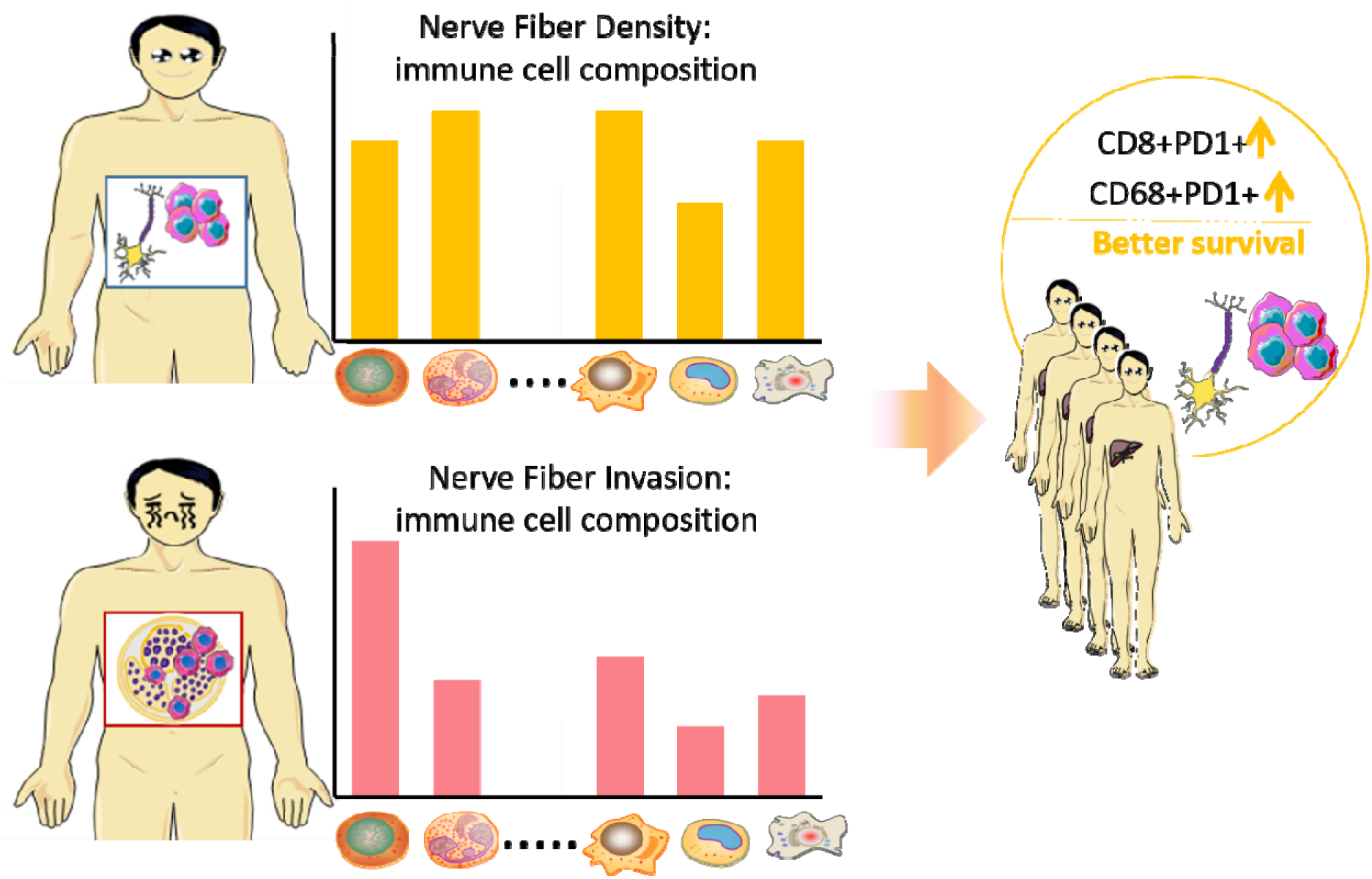

## Introduction

Cholangiocarcinoma (CCA) is a rare and challenging hepatobiliary malignancy arising from the biliary tract. Based on cancer location within the biliary tree, CCA is classified into 3 subtypes: intrahepatic cholangiocarcinoma (iCCA), perihilar cholangiocarcinoma (pCCA) and distal cholangiocarcinoma (dCCA). CCA usually has a 5-year overall survival (OS) of only <10%[1]. Up to now, surgical resection is still the only curative treatment for these patients, unfortunately with low percentages of patients being eligible for surgery. Patients are often diagnosed at an advanced stage and resection is no longer an option. For almost all CCA patients, conventional cytotoxic chemotherapy is the mainstay treatment option[2], providing only months of overall survival benefit and causing many toxic side-effects. The upcoming treatment options within personalized medicine has not brought much for the group of patients with perihilar cholangiocarcinoma[3]. Recent trials have opened the field of immunotherapy as a treatment option and if patients respond this has potential for a long term response[4]. For CCA patients this is a developing field and results from phase 3 trials are expected. Based on phase 1 clinical trials there is hope that immunotherapy in combination with chemotherapy regimens will improve outcomes in CCA patients as well[5].

The low success rate of CCA treatment is caused by many factors, and limited knowledge of its tumor microenvironment (TME) contributes to this problem. CCA has a high heterogeneity at the genomic, epigenetic and molecular level, hence, primary CCA contain a diverse range of cell types[6-11]. Also, the TME is host for many different immune cells and stimulatory and inhibitory effects take place. PCCA shows abundant desmoplastic stroma which contains many immune cells, providing either a host protective immune environment or tumor progression is facilitated[12]. Immune cell compositions play an important role in the immune response to the cancer and different phenotypes have been suggested in combined hepatocellular-cholangiocarcinoma patients (cHCC-CCA)[13] and in iCCA[14].

From a histopathological point of view, pCCA characteristically has an extensive stromal component and this is where the complex microenvironment interactions take place[12]. In the past decade, great efforts have been made to explore complexity of the TME and to develop novel therapies that might help to improve cancer patient outcomes. However, more needs to be discovered about the spatial relationship among cells within the complex TME and their expression patterns of co-stimulatory and inhibitory signals to understand the response to immune checkpoint blockades in clinical trials. Also, new biomarkers to predict response to checkpoint inhibitors are of urgent need, allowing selection of the right treatment for the patient.

With our latest studies, we have shown that nerve fiber density (NFD) in the TME functions as an important prognostic biomarker in CCA patients. NFD is associated with clinical outcomes in pCCA and iCCA patients[15, 16] and patients with presence of small nerve fibers in the TME show a better survival. The underlying mechanisms of this clinical observation are not discovered yet. Of main importance is the difference of a well-known aggressive feature perineural invasion or here named as nerve fiber invasion (NFI), which shows invasion of cancer cells into the nerve fibers (figure 1). This histological feature is detectable on a routine H&E staining and it is thought that the perineurium of the nerve fiber is a barrier for the chemotherapy to reach the cancer. Also, the nerve fiber environment provides a way of least resistance for the tumor to spread and progress. NFD has an opposite effect on outcome and consists of nerve fibers growing in the TME. In case of high NFD, small nerve fibers grow into the TME. These nerve fibers are only visualized by a special staining and the nerve fibers are smaller in diameter and usually don’t show any tumor invasion.

**Figure 1:**
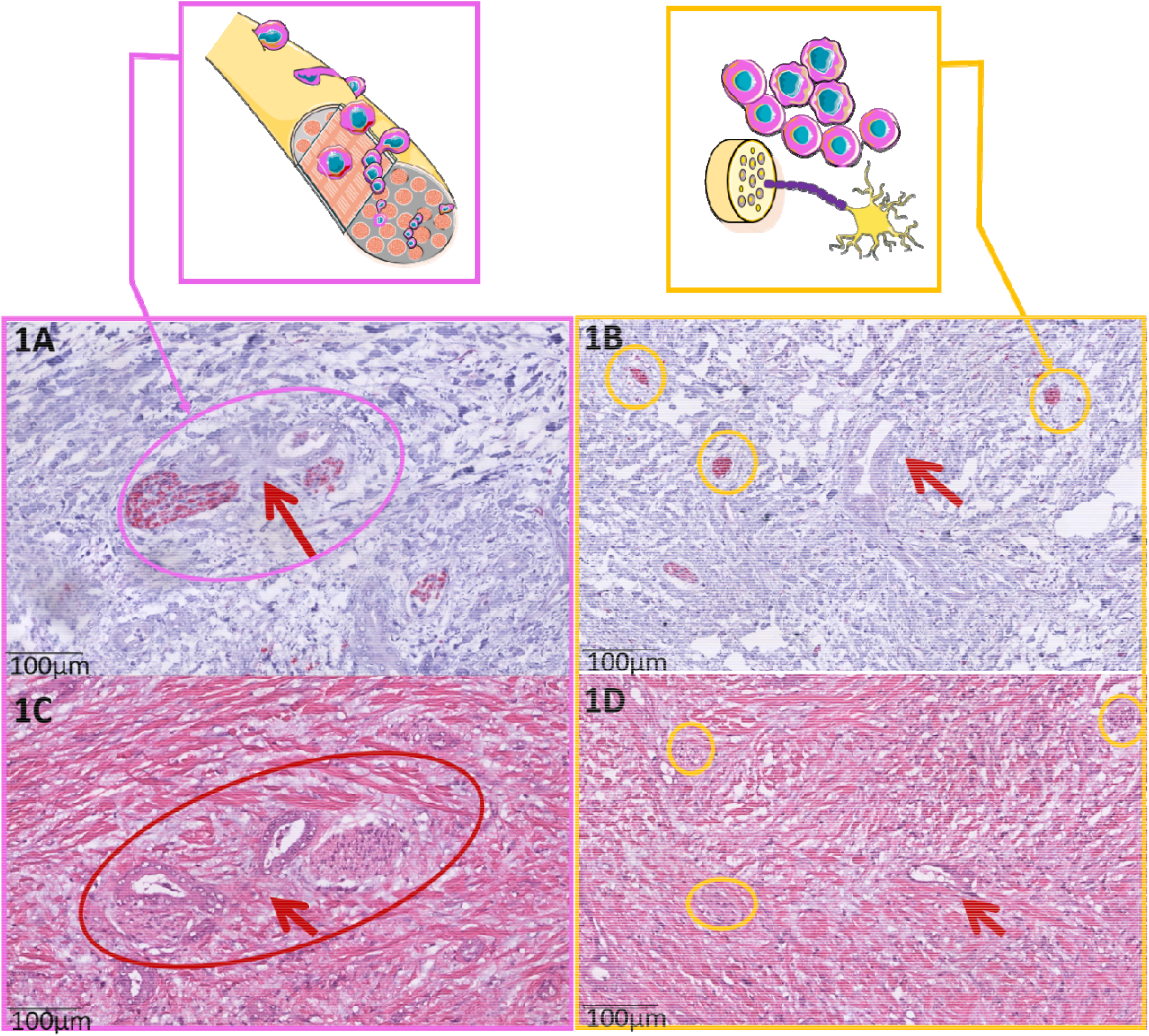
The difference between Nerve Fiber Invasion (NFI) and Nerve Fiber Density (NFD). 1A NFI is defined as tumor cells invading the perineurium of the nerve. In the neuronal marker (PGP9.5) you see the nerve fiber in red (red arrow) surrounded and invaded by tumor cells and glandular structures. 1B NFD shows the presence of small nerve fibers, found in the tumor microenvironment (TME). These nerve fibers are small in diameter (<10mu) and don’t show any invasion of tumor cells. The red arrow points to the tumor cells and the yellow circles mark the presence of small nerve fibers stained with the neuronal marker (PGP9.5). 1C Corresponding H&E staining of NFI. Nerve fiber invasion is recognizable for the pathologist. 1D H&E staining where the cancer is recognizable but the small nerve fibers are not detectable on this routine staining.

In this study, we use multiplex immunofluorescence (mIF) to reveal the differences in immune cell composition and distribution combined with the expression of co-stimulatory and co-inhibitory checkpoint markers between nerve fiber related phenotypes NFI and low vs high NFD in surgically resected pCCA patients.We hypothesized to see a different immunophenotype between the patients with NFI and high NFD since their OS is significantly different.

## Materials and Methods

### Patient Cohort

In total 47 pCCA formalin fixed paraffin embedded (FFPE) tissue blocks were selected from the archive of the University Hospital RWTH Aachen. All patients were treated and operated on in our hospital between 2010 and 2019. The study was conducted in accordance with the requirements of the Institutional Review Board of the RWTH-Aachen University (EK 106/18), the Declaration of Helsinki, and good clinical practice guidelines (ICH-GCP). Patient characteristics are provided in table 1.

**Table 1.**
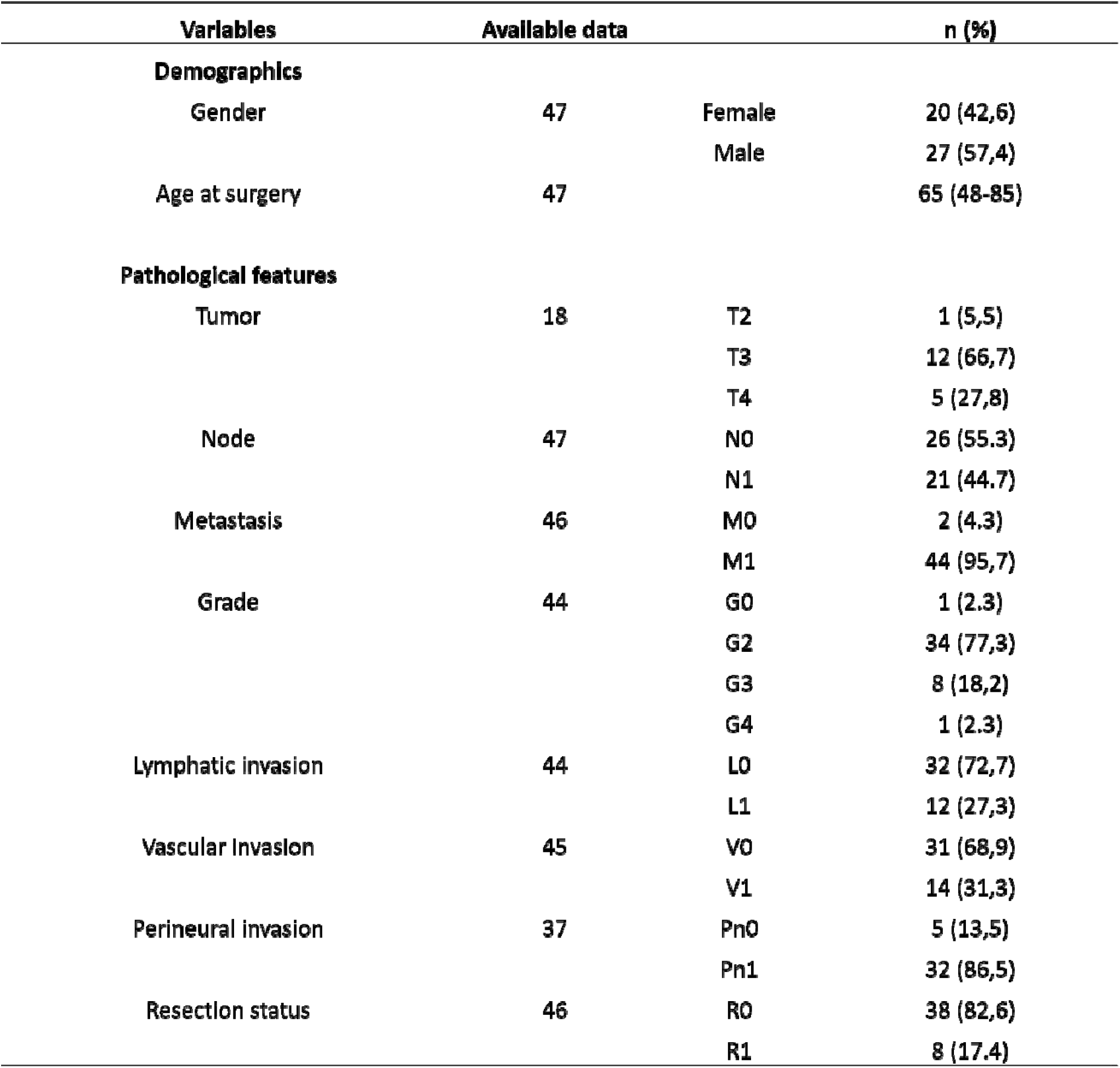
Patient characteristics.

| **Variables** | **Available data** |  | **n (%)** |
| --- | --- | --- | --- |
| **Demographics** |  |  |  |
| Gender | 47 | Female | 20 (42,6) |
|  |  | Male | 27 (57,4) |
| Age at surgery | 47 |  | 65 (48-85) |
| **Pathological features** |  |  |  |
| Tumor | 18 | T2 | 1 (5,5) |
|  |  | T3 | 12 (66,7) |
|  |  | T4 | 5 (27,8) |
| Node | 47 | N0 | 26 (55,3) |
|  |  | N1 | 21 (44,7) |
| Metastasis | 46 | M0 | 2 (4,3) |
|  |  | M1 | 44 (95,7) |
| Grade | 44 | G0 | 1 (2,3) |
|  |  | G2 | 34 (77,3) |
|  |  | G3 | 8 (18,2) |
|  |  | G4 | 1 (2,3) |
| Lymphatic invasion | 44 | L0 | 32 (72,7) |
|  |  | L1 | 12 (27,3) |
| Vascular Invasion | 45 | V0 | 31 (68,9) |
|  |  | V1 | 14 (31,3) |
| Perineural invasion | 37 | Pn0 | 5 (13,5) |
|  |  | Pn1 | 32 (86,5) |
| Resection status | 46 | R0 | 38 (82,6) |
|  |  | R1 | 8 (17,4) |

### Immunohistochemistry (IHC) in whole slide and image analysis

All samples were checked for the presence of tumor region by hematoxylin and eosin-stained sections. Slides were cut in tissue sections (2.5 μm thick) from formalin-fixed blocks, deparaffinized in xylene and rehydrated in graded alcohols. Slides were boiled in citrate buffer (pH 6.0) at 95–100°C for 5 min and were cooled for 20 min at room temperature with endogenous peroxide in methanol for 10 min. Then, these slides were incubated with rabbit anti-human PGP 9.5 (DAKO 1:100) overnight at 4°C to mark the nerve fibers. Histological Slides were scanned using the whole-slide scanners Aperio AT2 with ×40 objective (Leica Biosystems, Wetzlar, Germany), corresponding to a pixel-edge-length of = 0.252. A single digital image per case was uploaded in Qupath 0.1.6.

NFD nerve fiber counts from our previous study were used for immune cell phenotyping[16]. The NFD method was evaluated by manually counting the number of nerve fascicles with diameters of <100 μm in 20 continuous visual fields at ×200 magnification[17]. Based on NFD results, patients were categorized into a low NFD group (<10 nerve fibers) and a high NFD group (≥10 nerve fibers).

### Multiplex immunofluorescence staining

All FFPE samples were subjected to multiplex immunofluorescence (mIF) in serial 5,0 μm histological tumor sections obtained from representative FFPE tumor blocks which were confirmed with presence of the tumor region. The sections were labeled by using the Opal 7-Color fIHC Kit (PerkinElmer, Waltham, MA). The antibody fluorophores were grouped into a panel of 5 antibodies. The order of antibodies staining was always kept constant on all sections and sections were firstly counterstained with DAPI (Vector Laboratories). The multiplex immunofluorescence panel consisted of CD8, CD68, PD1, PD-L1, and PD-L2 (table 2). All antibodies were diluted with Antibody Diluent (with Background Reducing Components, Dako, Germany). Secondary antibodies were applied with ImmPRESS™ HRP (Peroxidase) Polymer Detection Kit (Vector Laboratories, US). TSA reagents were diluted with 1×□Plus Amplification Diluent (PerkinElmer/Akoya Biosciences, US).

**Table 2.**
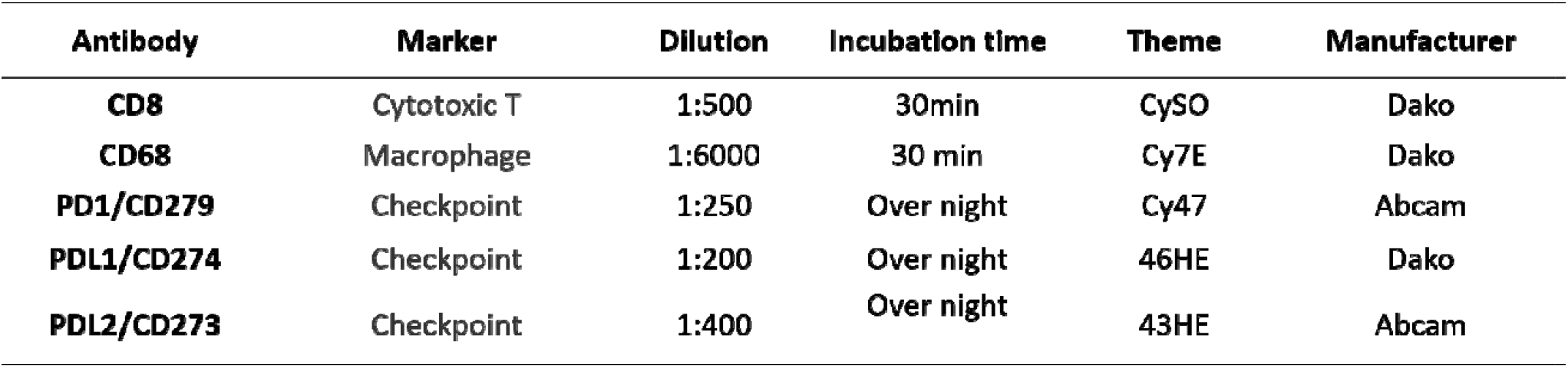
Monoclonal antibodies in panel 1.

The manual for mIF is described as Edwin R. Parra’s protocol[18]: in short, the first marker was incubated after the FFPE sections were deparaffinized in xylene and rehydrated in graded alcohols. The second marker was applied on the following day. And the third marker was applied on the third day. After all five sequential reactions, sections were finally cover-slipped with VECTRASHIELD® HardSet™ Antifade Mounting Medium.

The slides were then digitally scanned with the TissueFAXS PLUS system (TissueGnostics, Austria). Image analysis was performed in 3 regions of interest (ROI) in each image (only if present in the slide: tumor region, tumor free region, nerve fiber invasion region (NFI), the size of the ROI varying per slide. Immune cells were calculated in percentage throughout the whole project.

Strataquest software was used to analyse the antibody staining and cell counts. The library information was used to associate each fluorochrome component with a mIF marker. All immune cell populations were quantified as positive cells using the cell segmentation, thresholds were set manually under supervision of two pathologists (LH/MC). A positive cell was detected when expression exceeded the threshold value.

### Statistical analysis

All statistical calculations were implemented by using SPSS statistic software (v25, IBM, Armonk, NY).

Group comparisons were conducted by the Mann-Whitney U test or T-test in case of continuous variables, while the χ2 test, Fisher’s exact test in accordance with scale and number count were used in case of categorical variables. The Wilcoxon-matched-pairs-test or the paired t-test was applied to determine statistically significant differences between the different values, comparison was done between immune cells in different ROIs. Survival curves were generated by the Kaplan-Meier method and compared with the log-rank test. The level of significance was set to alpha= 0.05, and p values were calculated 2-sided testing. Overall survival (OS) was calculated as duration from the date of surgery to death of the patient. If patients were still alive or in case of a non-cancer related death, data was censored.

## Results

### Patients’ characteristics and ROIs

A total of 47 patients diagnosed with primary pCCA between 2010 and 2019 were included, 27 men (57.4%) and 20 women (42.6%) with a median age of 65 years (range 48-85 years) at surgery. All patients underwent surgical resection for pCCA. Clinicopathological details of the patients are outlined in table 1. After a follow-up until October 2021, the median OS for the whole cohort was 34 months (95% CI: 11–57).

For NFD: this study cohort was grouped into patients with a high NFD (n =13) and a low NFD (n =33) based on NFD analysis, already performed in previous work, which was performed on whole slide images using the Qupath digital software 0.1.6 version[19]. For NFI: miF digitized image analysis was performed based on tumor ROI (n =45), tumor free ROI (n =34) and NFI ROI (n =22) (see figure 2).

**Figure 2:**
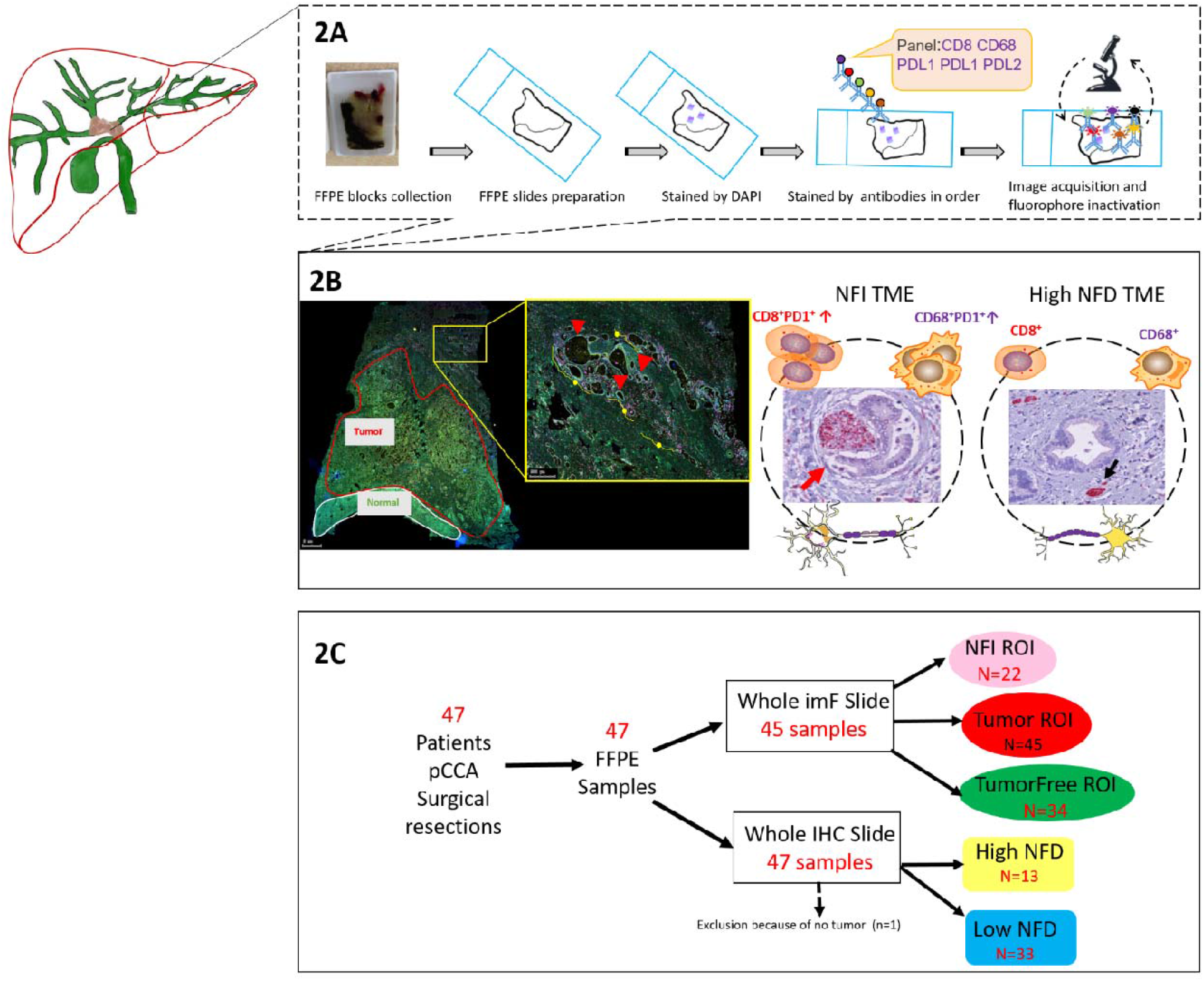
Overview of study workflow. 2A The formalin-fixed paraffin-embedded (FFPE) blocks were collected from the pathology archives. The slides were cut and stained with DAPI and 5 antibodies. The immunofluorescence stained slides were scanned. 2B The digital scans were annotated in different regions: Tumor, Tumor-free and NFI and cells were counted in these separate regions. For the NFD patients, the slides were selected based on the small nerve fiber count from previous work. For these patients the cells in the tumor region were counted. The tumor microenvironment (TME) was used for the positive cell counting. 2C In total we included 47 patients in this study. From 45 patients we were able to analyze the digital scans. Unfortunately not every slide contained all the regions. For example NFI was not detectable in each patient, the tumor free tissue was also not always available and sometimes the quality of the slide was not good enough to be analyzed (cracks or loss of tissue during antibody staining).

### Spatiality of immune cells in the TME of the NFI ROI

To specifically investigate the complexity and the phenotype of resident and infiltrating immune cells in the NFI region of pCCA, we applied 1 panel of 5-antibodies to phenotype the T-cells and macrophages in pCCA (see table 2 for an overview of the antibody panel). A full view of mIF image displays a complex and heterogeneous immune cell landscape in the TME. In short, cytotoxic T lymphocytes (CD8+ cytotoxic T cells) and CD68+ macrophages were measured in 3 ROIs (tumor, tumor free and NFI).

Analysis was performed on CD8+ T-cells and CD68+ macrophages in 3 ROIs (tumor, tumor free and NFI). This revealed higher numbers of CD8+ T-cells in the tumor ROI compared to the tumor free ROI (Wilcoxon matched pairs test, p= 0.027) in pCCA. For CD68+ macrophages there were significant differences between tumor free ROI and tumor ROI, with higher numbers of CD68 macrophages in the tumor ROI (Wilcoxon matched pairs test, p=<0.001). Also the NFI ROI showed even higher numbers of CD68+ macrophages compared to the tumor ROI (Wilcoxon matched pairs test, p= 0.034) and to the tumor free ROI (Wilcoxon matched pairs test, p= 0.016) in pCCA patients (figure 3).

**Figure 3:**
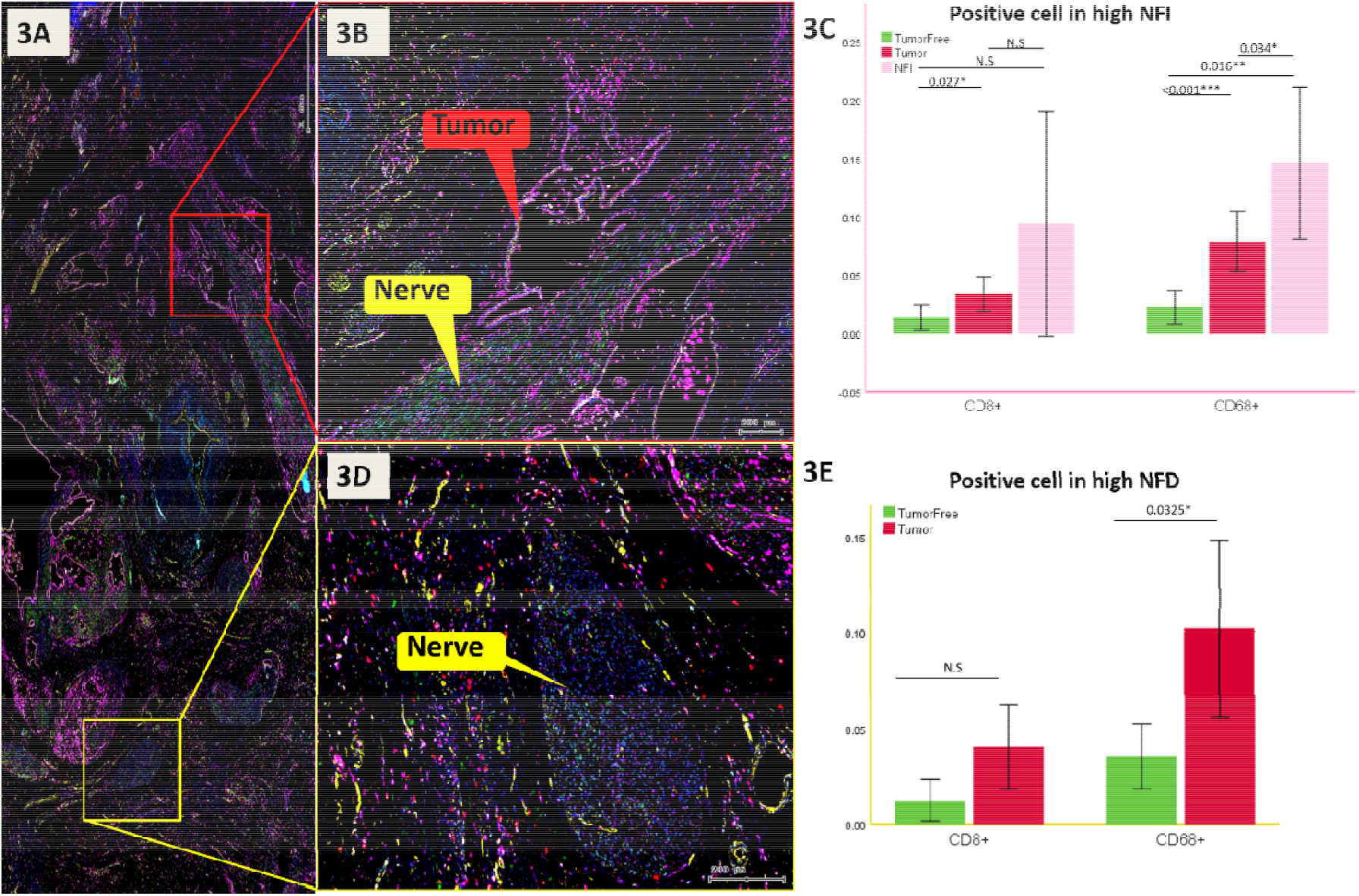
multiplex immunoflorenscence (mIF) digitized images. 3A Zoomed-in image of a perihilar cholangiocarcinoma, with the presence of many big nerve fibers and large vessels in the hilusregion of the liver. 3B The red box visualizes NFI with the tumor glands highlighted with red and the nerve fiber with tumor invasion highlighted in yellow. The tumor glands are orientated really close to the nerve and invade the perineurium. 3C For NFI the CD8 cell counts were significantly higher in the tumor region compared to the tumor free region (p= 0.027). CD68 was also more expressed in the tumor region compared to the tumor free region (p= 0.001), the NFI region showed significantly more expression of CD68 compared to the tumor region (p= 0.034) and tumor free region (p= 0.016). 3D The yellow box visualizes a large nerve fiber without tumor invasion. The increase of small nerve fiber which are counted for a high NFD are not detectable without a neuronal marker. The positive cell counting was done in the tumor annotation in the patients proven to have a high NFD based on previous work. 3E For NFD positive cell counts for CD8 were not significant between the tumor region and the tumor free region. For CD68 there were significantly more macrophages in the tumor region compared to the tumor free region (p= 0.0325).

### Survival analysis of patients with NFI, low and high NFD

OS survival analysis between the subgroups showed a significant difference between the group high NFD vs NFI (p= 0.032) and high NFD vs low NFD (p= 0.046). The median OS of high NFD patients was 92 months (95% CI:41-142) compared to a median of 20 months (95% CI: 4-35) in patients with a low NFD (p= 0.046) and compared to the median OS of patients with NFI of 19 months (95% CI: 7-33) (p= 0.032).

We compared OS for the NFI group of patients grouped in low and high CD8 counts. The OS shows a significant better OS with a median OS of 74 months (95% CI: 54-93) in the patients with high CD8 counts compared to a median OS of 48 months (95% CI: 17-78) in patients with a low CD8 counts (p= 0.044). Then, OS for the NFD group of patients divided in low and high CD8 counts, didn’t show significance in the OS (p= 0.091).

When we compared OS for the total group of patients (n=45 patients) with CD8+PD1+ expression grouped in low (n=30) and high (n=15) CD8 counts combined with PD1 expression there was a significant difference in OS with a median of 92 months OS (95% CI:39-145) in patients with high CD8+PD1+ expression compared to a median of 25 months (95% CI: 10-40)) in patients with a low CD8+PD1+ expression (p= 0.03).

For OS for the total group of patients (n=45 patients) with CD68+PD1+ expression, grouped in low (n=18) and high (n=27). CD68 counts combined with PD1 expression showed a significant difference in OS with a median of 55 months (95% CI:35-74) in patients with high CD68+PD1+ expression compared to a median of 20 months (95% CI: 5-35) in patients with a low CD68+PD1+ expression (p= 0.04). (Figure 4 for Kaplan Meier curves).

**Figure 4:**
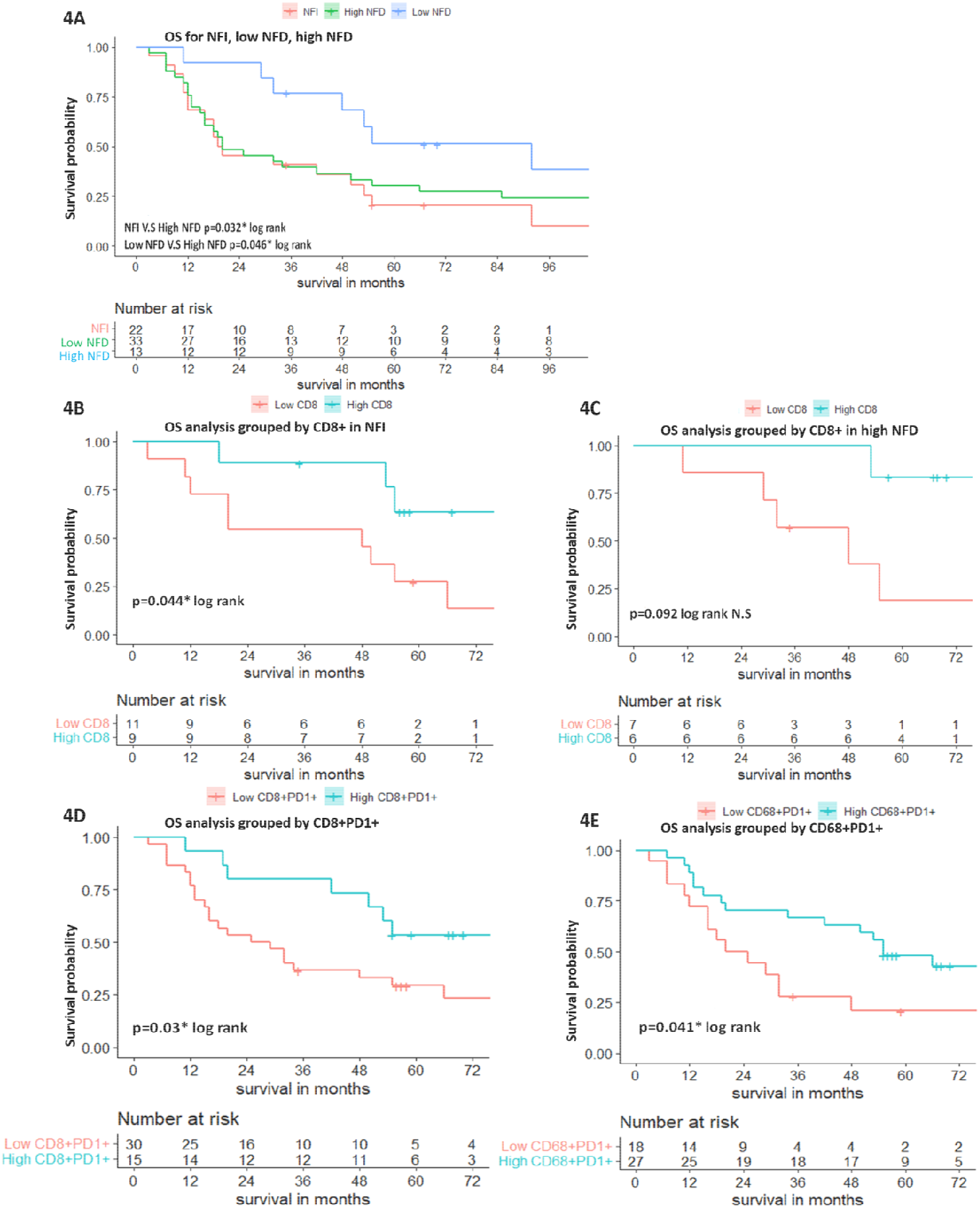
Survival analysis presented in Kaplan Meier Curves. 4A Oncological survival in pCCA patients, providing an overview NFI (n= 22), low NFD (n= 33) and high NFD (n=13) groups. The median overall survival (OS) of high NFD patients showed 92 months (95% CI:41-142) compared to a median of 20 months (95% CI: 4-35)) in patients with a low NFD (p= 0.046) and compared to the median OS of patients with NFI of 19 months (95% CI: 7-33) (p= 0.032). 4B OS for the NFI group of patients grouped in low and high CD8 counts. The OS shows a significant better OS with a median OS of 74 months (95% CI: 54-93) in the patients with high CD8 counts compared to a median OS of 48 months (95% CI: 17-78) in patients with a low CD8 counts (p= 0.044). 4C. OS for the NFD group of patients in low and high CD8 counts, didn’t show significance in the OS (p= 0.092). 4D OS for the total group of patients (n=45 patients) with CD8+PD1+ expression, grouped in low (n=30) and high (n=15) CD8 counts combined with PD1 expression showed a significant difference in OS with a median of 92 months OS (95% CI:39-145) in patients with high CD8+PD1+ expression compared to a median of 25 months (95% CI: 10-40)) in patients with a low CD8+PD1+ expression (p= 0.03). 4E OS for the total group of patients (n=45 patients) with CD68+PD1+ expression, grouped in low (n=18) and high (n=27). CD68 counts combined with PD1 expression showed a significant difference in OS with a median of 82 months (95% CI:35-74) in patients with high CD68+PD1+ expression compared to a median of 20 months (95% CI: 5-34) in patients with a low CD68+PD1+ expression (p= 0.041).

These data indicated that pCCA patients with high PD1 expression in the tumor have a better OS survival.

### Positive cell comparison between NFI ROI and NFD in patients with pCCA

We investigated NFD in association with CD8^+^ T-cells and CD68^+^ macrophages as well as their co-inhibitory and co-stimulatory checkpoint markers which were reported to play a critical role in the process of immune escape mechanisms. We hypothesized that the reason for the better prognosis of high NFD patients was to be found in differences in the immune cell composition and spatiality. The patients with a NFI ROI (n=20) were analyzed and positive cell counting per region of ROI was performed (figure 5).

**Figure 5:**
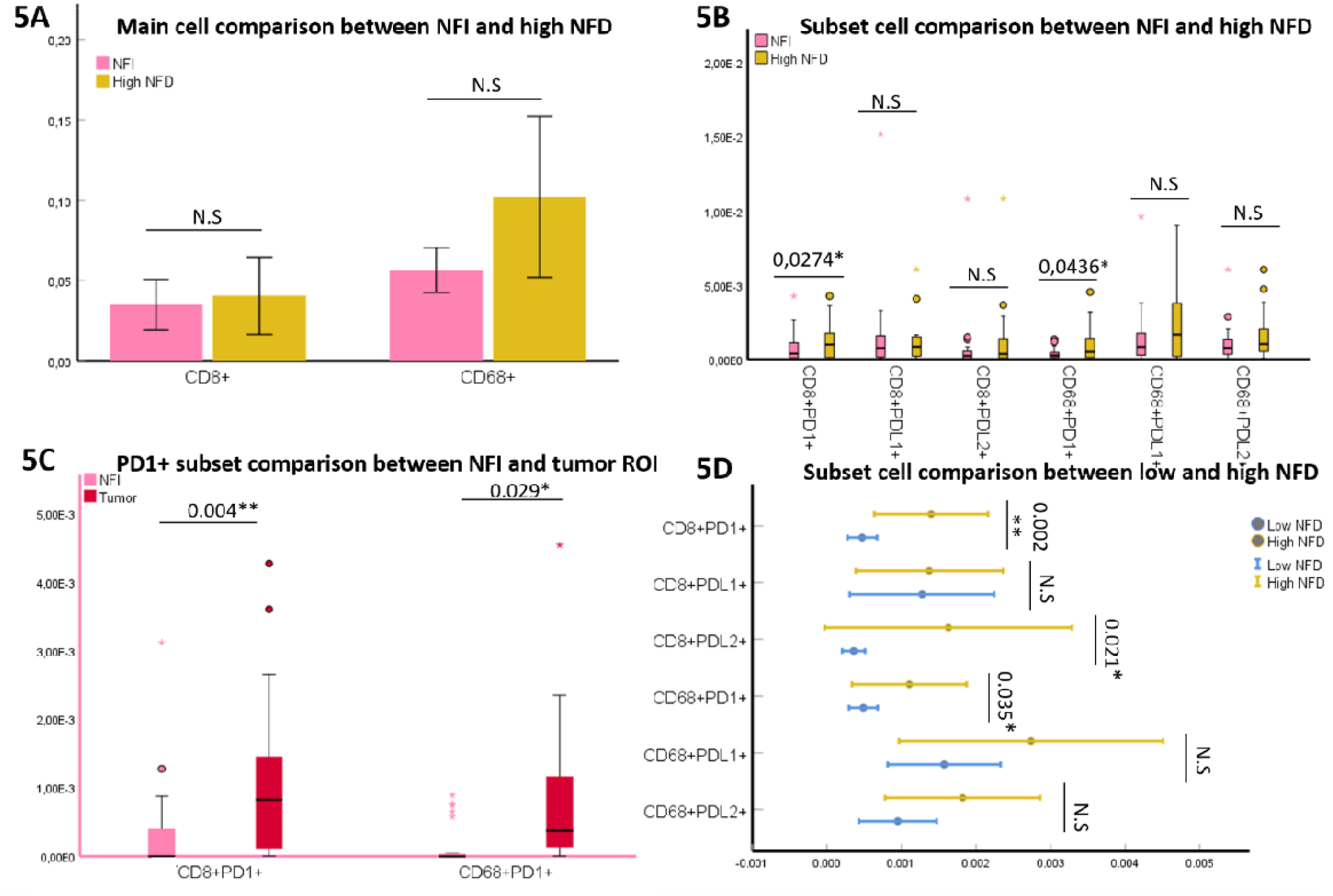
Cell comparison between different ROIs and low vs high NFD. 5A Main cell comparison between NFI and high NFD patients. The CD8+ and CD68 cell count didn’t show a significant difference. 5B Subset cell comparison between NFI and high NFD patients. High NFD patients showed significant higher expression of CD8+PD1 and CD68PD1 (p= 0.0274 and p= 0.0436 respectively). 5C Subset analysis between NFI and tumor ROI. The tumor ROI showed significant higher levels of CD8+PD1 and CD68PD1 cells (p= 0.004 and p= 0.029 respectively). 5D Subset cell analysis between the expression of PD1, PD-L1 and PD-L2 in patients with a low and high NFD. Patients with high NFD compared to patients with a low NFD showed significant higher levels of expression in CD8+PD1 (p= 0.002), CD8+PD-L2 (p= 0.021) and CD68PD1 (p= 0.035). The expression of CD8+PD-L1, CD68PD-L1 and CD68PD-L2 didn’t reach significance.

Here, we observed no significant difference between the CD8+ T-cells and CD68+ macrophages in the NFI ROI and high NFD.

### Expression of co-stimulatory and co-inhibitory receptors and ligands

We found that signals of co-expression of CD8+PD1+ as well as CD68+PD1+ were higher in the high NFD patients (n=13) when compared to the NFI ROI (n=20) (Mann-Whitney U-test; p= 0.0274, p= 0.0436, respectively, figure 5B). Next, we examined the expression of the co-inhibitory receptor PD1 between tumor and NFI ROIs through paired comparison, compared-sample T-test. This showed a significance between CD8+PD1+ and CD68+PD1+ in the tumor and NFI ROIs (p= 0.004, and p= 0.029, respectively, figure 5C).

The significant distribution of co-stimulatory and co-inhibitory receptors are illustrated in figure 5D, CD8+PD1+, CD8+PD-L2+, and CD68+PD1+ were all abundant in high NFD patients when compared to low NFD patients (Mann-Whitney U-test; p= 0.002, p= 0.021, p= 0.035, respectively). This low expression of PD1 in the NFI region, suggests that NFI might be a region difficult to reach for immunotherapy effects.

## Discussion

PCCA is considered a rare primary biliary tract cancer and therefore it remains understudied. Large cohorts of patients are lacking and patients usually have a poor prognosis, especially those who are not eligible for a surgical resection.

Immunotherapy doesn’t have a big place yet in the treatment of pCCA and stratification of patients for the right adjuvant treatment still needs to be improved. For other liver cancers, as HCC, immunotherapy in combination with Bevacizumab is already a first line treatment[20]. First clinical trial results report that pCCA is a immunoresponsive malignancy indicating a potential role in improving patients survival. Still low numbers of patients respond to immunotherapy and biomarkers to stratify patients for immunotherapy are of urgent need[21].

The histology of pCCA usually shows characteristic growth pattern of nerves invaded by cancer cells. Even though most patients have this feature somewhere present in the tumor, not all patients fare equally poor. In this study we have shown that NFD and PD1 are co-expressed in patients with a significantly better survival. The underlying mechanism for this still needs to be further investigated.

NFD is defined as large numbers of small nerve fibers in the TME, these nerve fibers are not invaded by cancer cells. A latest study has demonstrated that CD8+ infiltration was associated with better survival in patients with iCCA[22]. Hence, we evaluated the clinical significance of the main immune cells (T cells and macrophages) in pCCA patients. In our previous publication[16], we reported the role of NFD in pCCA and presented a positive correlation with high NFD being independently associated with better survival after surgical resection. We hypothesized that the small nerve fibers attract immune cells providing a better immune response to the cancer. Patients with a high NFD have abundant CD8+PD1+ T-cells and CD68+PD1+ macrophages.

This finding identifies a subgroup of pCCA patients with a better survival, making them eligible for adjuvant treatment in terms to reach a longstanding response.

PD1 checkpoint therapy unleashes the immune cells which expands the T cell population at the interface and in the tumor. Potentially the high NFD subgroup of patients could benefit from immunotherapy, when the CD8+ T-cells blocked with PD1 are released. Our data are in line with previous findings on HCC patients, where patients with high levels of PD1 expression showed an improved survival[23] and low counts of CD8 T-cells are indicative for a poorer outcome[24]. Previous work on the immune landscape in intrahepatic cholangiocarcinoma showed an immunosuppressive environment with low numbers of CD3 and CD4 was correlated with an poor outcome[25] and low expression of PD1 correlated with a better outcome[26-28]. For extrahepatic cholangiocarcinoma high numbers of CD3+ T-cells combined with expression of PD-L1 on the tumor cells was correlated with a more invasive growth[29]. The prognostic relevance of the PD1 marker is therefore diverse but expression of this checkpoint receptor indicates patients are likely to benefit from immunotherapy[30-32]. Previous work has shown that cholangiocarcinoma patients with high densities of TILs also have high expression levels of PD-L1[33]. Besides using the expression of PD1 and PD-L1 to predict outcome, different immune responses in biliary tract cancer are indicative for a better or worse outcome[34]. The different prognostic values of the PD1 and PD-L1 expression in cholangiocarcinoma patients suggests that expression of this marker by itself is not enough to function as a good biomarker. On the other hand we have investigated the NFI ROI and illustrated a different immune cell signature with different expression levels of the co-stimulatory and co-inhibitory markers. This is of relevance while the nerve fiber invasion is thought to be a route for the tumor to progress. The invasion of tumor cells into the perineurium provides a protective environment for the cancer cells. Supporting this hypothesis the perineural invasion is often found at distant sites from the main tumor bulk and can be the reason for a non-in-sano resection.

Also it is described that the perineurium is a barrier for therapeutic agents to reach and response to neoadjuvant treatment is not often observed in the nerve fiber invasion region. For future developments of therapeutic approach it is important to consider this phenomenon as an obstacle in reaching a complete response of the cancer, since our data show that PD1 is not expressed specifically in this region.

The strategy to treat the cancer with therapeutic agents possibly contains a combination therapy, including immunotherapy and agents that targets the nerve fiber invasion ROI. Here we show that the NFI ROI shows a comparable immune cell composition as the tumor ROI but the expression of checkpoint signals is significantly different and PD1 levels are significantly lower at this site. This is an important observation for future immunotherapy because the effect of PD1 blockers is likely to be less at the sites showing nerve fiber invasion.

To our knowledge, this is a first study in pCCA using a wide multiplex antibody panel focusing on immune cells in the TME in combination with checkpoint markers correlated with nerve fiber density and nerve fiber invasion. PD1 expression correlates with high NFD patients suggesting NFD as a simple prognostic biomarker. NFD can be easily integrated in the routine workup of the pathology report, since only one neuronal antibody is needed to achieve a nerve fiber count.

## List of abbreviations

CCA: Cholangiocarcinoma
CHCC-CCA: Combined hepatocellular-cholangiocarcinoma
DCCA: Distal cholangiocarcinoma
FFPE: Formalin fixed paraffin embedded
HCC: Hepatocellular carcinoma
ICCA: Intrahepatic cholangiocarcinoma
MIF: Multiplex immunofluorenscence
NFD: Nerve fiber density
NFI: Nerve fiber invasion
OS: Overall survival
PCCA: Perihilar cholangiocarcinoma
PD1: Programmed Death receptor
PD-L1: Programmed Death Ligand 1
PD-L2: Programmed Death Ligand 2
PGP 9.5: Protein gene-product 9.5
ROI: Region of interest
TME: Tumor microenvironment

